# PEMPS: A Phylogenetic Software Tool to Model the Evolution of Metabolic Pathways

**DOI:** 10.1101/2024.01.04.574206

**Authors:** Nicholas S. McCloskey, Ayna Mammedova, David A. Liberles

## Abstract

**Background:** Metabolic pathways support the enzyme flux that converts input chemicals into energy and cellular building blocks. With a constant rate of input, steady-state flux is achieved when metabolite concentrations and reaction rates remain constant over time. Individual genes undergo mutation, while selection acts on higher level functions of the pathway, such as steady-state flux where applicable. Modeling the evolution of metabolic pathways through mechanistic sets of ordinary differential equations is a piece of the genotype-phenotype map model for interpreting genetic variation and inter-specific differences. Such models can generate distinct compensatory changes and adaptive changes from directional selection, indicating single nucleotide polymorphisms and fixed differences that could affect phenotype. If used for inference, this would ultimately enable detection of selection on metabolic pathways as well as inference of ancestral states for metabolic pathway function.

**Results:** A software tool for simulating the evolution of metabolic pathways based upon underlying biochemistry, phylogenetics, and evolutionary considerations is presented. The Python program, Phylogenetic Evolution of Metabolic Pathway Simulator (PEMPS), implements a mutation-selection framework to simulate the evolution of the pathway over a phylogeny by interfacing with COPASI to calculate the steady-state flux of the metabolic network, introducing mutations as alterations in parameter values according to a model, and calculating a fitness score and corresponding probability of fixation based on the change in steady-state flux value(s). Results from simulations are consistent with *a priori* expectations of fixation probabilities and systematic change in model parameters.

**Conclusions:** The PEMPS program simulates the evolution of a metabolic pathway with a mutation-selection modeling framework based on criteria like steady-state flux that is designed to work with SBML-formatted kinetic models, and Newick-formatted phylogenetic trees. The Python software is run on the Linux command line and is available at https://github.com/nmccloskey/PEMPS.

## Background

Metabolic pathways support the enzyme flux that generates energy and building blocks to sustain life [1]. Pathways evolve with constraints on functions at the pathway level driven by mutations occurring at the underlying level of individual enzyme properties [2]. Selection can also act on different levels of pathway function, including time-dependent dynamics and steady-state flux [2]. Studying the effect of selective pressure on a metabolic network’s steady-state flux, a condition achieved when metabolite concentrations and reaction rates are constant in the network [3], can elucidate its co-evolutionary dynamics [4]. Modeling the evolution of metabolic pathways is a piece of the larger genotype-phenotype map [2]. If used for inference (by using methods like Approximate Bayesian Computation [5] that use simulation for inference), such models have the potential to detect single nucleotide polymorphisms and fixed differences that could affect phenotype, ultimately enabling detection of lineage-specific selection on metabolic pathways as well as inference of ancestral states at phylogenetic tree nodes for metabolic pathway function [2]. The first step in developing such an inference tool is building a model for simulation.

Because most genes function with epistatic effects on other genes involved in the same network or pathway, the phenotypic effect of their mutation could depend on biochemical context, something lost in polygenic risk scores in statistical genetics that assumes that every gene functions independently [2]. The rules for dependent evolution are governed by mathematical expressions known to biochemists that obviate the need for mechanism-free statistical genetics approaches [2]. Under mutation-selection-drift balance, rapid evolution of individual enzymes can occur without altering pathway flux, due to intermolecular epistasis in which the functional change in one enzyme affects selective constraints and evolutionary rates of other enzymes in the network [4]. Such evolution in multiple enzymes could be due to directional selection on intermediates or to compensatory changes, but a biochemically informed null model of pathway evolution assuming stabilizing selection is required to make this distinction. The PEMPS software tool is presented to address the need for a framework that incorporates epistasis in the systemic context of metabolic pathway evolution across species, as opposed to studying the evolution of individual network components assuming their evolutionary independence. While metabolic pathway evolution simulations [6–8] – even over a phylogeny [9] – have been conducted for various studies, research software for simulation using an input metabolic model and phylogeny had previously been unavailable.

Three big-picture hypotheses of pathway regulatory evolution have been proposed. First is the topological selection hypothesis, based on phenomenological and mechanistic models which posits that selection on pathway flux control is constant, driven by strong selection early in pathways and on committed steps with large kinetic or thermodynamic barriers between substrates and products [10]. Second, the mutation-selection-drift balance hypothesis is based on theoretical work demonstrating that intermolecular epistasis can preserve pathway flux while its individual enzymes evolve rapidly, and posits that changes in enzyme kinetic properties cause a shift of flux control over evolutionary time due to mutation-driven processes [11, 12]. Third, the frequent soft sweeps hypothesis proposes that flux control shifts rapidly due to balancing selection on seasonal timescales with action occurring on standing variation without a role for new mutation [13]. The software presented here is designed to simulate under the second hypothesis, but could in principle be tuned to simulate under the alternative hypotheses as well. A simple shift in the distribution of mutational effects from one dominated by slightly deleterious mutation to one dominated by neutral mutation with equal frequencies of deleterious and adaptive changes would give rise to dynamics consistent with the first rather than the second hypothesis [11] but is not expected biologically [14].

Among the possible descriptions of a metabolic pathway, kinetic modeling offers the most detail. In this framework, each enzymatic step in the network can be represented with an enzyme-kinetic rate law such as the Michaelis-Menten equation [3]. This equation is a function of enzyme concentration, substrate concentration, catalytic rate constant, and Michaelis binding constants, and describes the enzyme-substrate complex as an equilibrium process [15]. The change in concentrations of metabolites over time is determined by the sum of the rates synthesizing the metabolite minus the sum of the rates consuming it. After accounting for all metabolites in the pathway, this results in a system of ordinary differential equations (ODEs) whose solution represents the metabolite concentrations required for the network to reach steady state. Such deterministic kinetic models based on systems of ODEs have been the most frequently used approach to metabolic modeling [3]. Models can behave differently depending on their parameters. Some reach steady state, where all fluxes balance each other such that metabolite concentrations do not change. Alternatively, the system can exhibit oscillatory behavior, or the concentrations could all drop to zero. The probability of oscillation or other instability has been shown to increase with model complexity; for large kinetic models, it is unusual to reach steady state, and this may have been selected for [3].

Haldane’s relationship dictates that the dynamic equilibrium parameters in an enzyme reaction are not free to vary independently, but that any mutations must comply with the biophysical (energetic) constraints of the reaction landscape. This is embedded into the profiles of linked reaction parameters in the mutational profile [14]. The previous study demonstrated that fitness equilibrium is reached whenever selection is present, but that some parameters (binding constants) can continue to show systematic directional movement without affecting fitness [2].

Fitness effects of new mutations have been shown to have large fractions of both severely (i.e., lethally) and slightly deleterious mutations, with the magnitude and frequency of such effects depending on the mutational landscape of the coding sequence [2]. A mutation-selection model (see Methods) treats mutational and fixation probabilities independently [16], and unlike a Wright-Fischer approach can accommodate large population sizes without incurring a prohibitive computational cost by assuming that one mutation fixes before the next one is introduced and eliminating the need to model the entire population. A tool to simulate the evolution of metabolic pathways and their regulation was generated using this framework.

### Implementation

PEMPS is a Linux command line-executable Python program that can simulate the evolution of an input metabolic pathway over an input phylogeny and report the changes in model parameters along the lineages of the phylogenetic tree over evolutionary time. The user provides commands in a text file designating the underlying biology, including a metabolic model, the reaction flux(es) on which selection operates and the relative weights in the calculation of selective coefficients, a phylogeny as a Newick string for the species, the ploidy of the genome, population size(s) of all branches, any non-genetic parameters to be excluded from mutation, and the number of simulations to run. A future release could enable lineage-specific shifts in the optimal flux for a pathway. The user is given control over the network dynamics, as they determine which model parameters are held constant and on which reaction fluxes selection acts. A different decision about the target of selective pressure results in new evolutionary constraints on different constellations of model parameters in the network, which is likely to affect the changes observed in parameters over phylogenetic time. A flowchart of the overall program function is presented in Figure 1.

**Figure 1.**
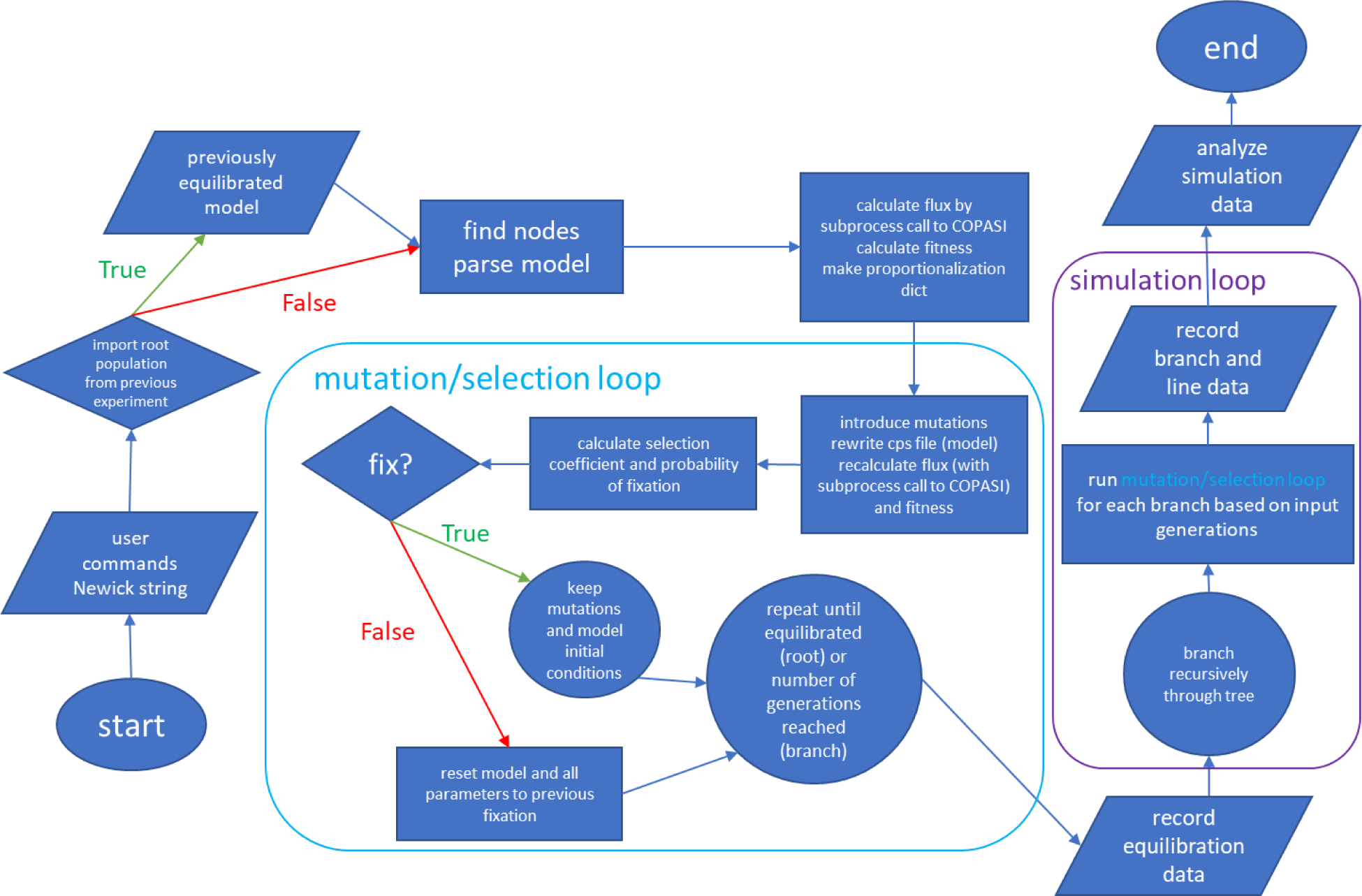
PEMPS program flowchart.

The program requires a Newick string from the user-specified file for which the branch simulation proceeds recursively. The tree must be rooted [17]. Rooted trees can be obtained from the NCBI taxonomy [18], TimeTree [19], rooted using gene tree/species tree reconciliation with software like SoftParsMap [20] or NOTUNG [21], or by software like DendroPy [17] using techniques like mid-point rooting when species relationships are unknown [22]. PEMPS requires branch lengths as substitutions per site or in millions of years to determine the number of generations to simulate for each branch. The input metabolic model is prepared with the GUI version of COPASI, and the SE version, interfaced through Python, is used to calculate steady-state flux. As the fitness calculation in the mutation-selection process depends on steady-state flux (where selection is based upon this criterion), the program is only compatible with models that can reach steady state with their original parameters. Detection of binding constants for mutation and Haldane’s constraint depends on informative parameter naming, which some models do not provide (the label could be *parameter 12* as opposed to *KmATP*). In such cases, the user would have to offer specifications in the commands file.

The program conducts a forward-time simulation with discrete generations using a mutation-selection model [23]. In each generation, each parameter has a small probability (Poisson random variable, lambda = 0.003) of mutating, and if mutation occurs, the COPASI file is rewritten for the recalculation of steady-state flux with the mutated parameters. To model the tendency towards slightly deleterious change, mutational effects are drawn from a normal distribution centered at −1% for all parameters except binding constants (1%), because poorer binding corresponds to a larger Km value. The assumption that mutations tend towards slight functional degradation is consistent with results from mutation accumulation experiments [12] and parameterizations of the distribution of fitness effects in protein coding genes [14, 24]

The user can hold any global or kinetic parameters to their starting values by prohibiting their mutation, which could be useful, for example, to incorporate an assumption of constantly replenishing supply of the network’s input(s). Most metabolic models explored during development of the program are constructed this way, and without any specification, the network inputs remain constant throughout the simulation. To implement Haldane’s constraint, the program collects each reaction’s maximum velocity, equilibrium constant, and binding constants, provided that they are informatively named (which tends to be the case) and that there are equal numbers of substrates and products, so that pairings can be established. After a mutation to either a forward or reverse binding constant, the reverse reaction velocity is calculated, and the counterpart parameter is updated accordingly. An analogous procedure would be applied to the catalytic constant and its reverse, although most models are not specified with reverse kcat values.

To model selective pressure on the steady-state flux of a subset of the network’s reactions, population fitness is calculated from proportionalized reaction flux (x) with the logistic function given in equation 2.

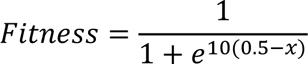

A logistic curve is suitable because the upper asymptote represents a diminishing marginal return that corresponds to the biomolecular reality that although a cell cannot survive with zero flux, increasing production beyond a certain threshold results in no further fitness advantage. Some contexts, however, where there is effectively no ceiling to the benefit of increasing cellular product would be best modeled with a linear function, as might be the case for the generation of materials required for cell division during the exponential growth phase in bacteria. Because multiple reaction fluxes may be under selection, and these values may lie on different scales, the fluxes are proportionalized based on initial (pre-mutation) flux values such that the starting fitness score for each flux is 1. Fitness scores are calculated for each of these proportionalized flux values and adjusted such that a minimum fitness of 0 corresponds to no enzyme flux, a fitness of 1 corresponds to the starting flux, and the asymptote represents the production threshold beyond which no further fitness advantage is gained. The population fitness is then the weighted geometric mean of these normalized scores. This function can be modified in the code by a user.

The population fitness values from before and after a mutation are used to calculate the probability of fixation with Kimura’s formula (Equation 3) [25] as presented by Otto and Whitlock [26].

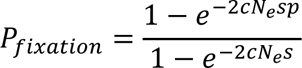

The selection coefficient *s* is the ratio of the new population fitness (after mutation) to the old fitness minus 1. The initial frequency parameter p is usually calculated by 1/(*cNe*) where *c* is the ploidy and *Ne* is the effective population size, but to speed up simulation, this parameter is set to 0.5 [27]. This gives neutral mutations a 50% chance of fixing, accelerating the neutral walk across sequence space.

Before the simulation runs through the tree, the “root” population undergoes mutation and selection until its fitness and most (80%) of its reaction fluxes reach values that remain stable over many generations. This state of equilibrium is determined by calculating the slope of the line of best fit and coefficient of variation (values indicating “flatness” of the line) of the select parameters over 100-fixation windows. When the values of five consecutive windows are within certain thresholds of zero for fitness and the majority of fluxes, equilibration continues for as many generations as had transpired by that point, and then branching simulation commences. Because equilibration occurs before branching, the phylogeny is not relevant in this step, which means that if the starting population size, ploidy, and fluxes under selection are the same, one equilibrated population can act as the starting point for any other tree the user wants to simulate with the same starting population parameters and metabolic model. The user has the option to import an equilibrated set of parameters into the root node population and can either equilibrate further or start the phylogenetic simulation.

Branch lengths as either millions of years or substitutions per site are required, and the number of generations to be simulated for each branch depends on the length and user-specified measurement type. To convert branch lengths to number of generations to simulate, the program determines at what number of generations during equilibration an average number of 7 mutational proposals are introduced to both enzyme concentration and enzyme functional parameters across the model reactions. This threshold is based on the following assumptions: (1) the two-percent rule which states that branching avian and mammalian lineages tend to differ from each other by 1% every 1 million years [28]; (2) that 50% of neutral changes fix; (3) a bias towards deleterious change; (4) that the coding sequence for a typical enzyme contains ∼250 amino acids subject to the mutational effect distribution for enzyme parameters and ∼250 sites that can affect gene expression leading to changes in protein concentration. This latter category would include mutations in promoter sites that influence transcription factor binding [29], which affect the quantity of transcripts and ultimately proteins produced, thus causing changes in the concentration of enzyme ([E]). In models parameterizing [E], PEMPS probabilistically proposes mutations to [E] and other enzyme parameters in a 1:1 ratio. If the input branch lengths are measured in millions of years, this threshold is simply multiplied by branch length to calculate simulated generations per branch; if measured in substitutions per site, a factor of 100 is introduced to convert branch lengths to percentages.

Lineages (edges) are formed by connecting branch simulations when tracing a path directly from the equilibrated root node through all internal nodes to an external node, with bifurcating branches each starting their separate simulations with the same set of inherited parameters. Larger branch lengths result in a higher number of generations along the lineage, affording more opportunity to fix new mutations, which creates an expectation of greater change along longer lineages. Population sizes are also determined by the user, who can choose to set a uniform value across the entire tree or enter the population size individually for each branch. A new equilibrium may not be reached after branching to a new population size by the next bifurcation, but this corresponds to real evolution. Output generated by the program includes the changes in parameters for all mutations, fixed and unfixed, in tabular format, graphs of parameters for lineage and branch, and heatmaps of percent differences from starting values for lineages and branches (more detailed description of output can be found in the program docstrings (https://github.com/nmccloskey/PEMPS). When multiple simulations are run (the user’s decision), tables of average percent differences for each parameter for lineages and branches are produced with corresponding heatmaps.

## Results

Data from three example runs of the program are presented. The first network is a model of the pyruvate branches in the bacterium *Lactococcus lactis* [30]. The following increasingly complex models based on glycolysis [31, 32] were selected because this pathway is found with conserved function across the tree of life and has been well studied [4, 33]. Furthermore, it has been shown that the structure of the enzymes involved changes across species, and variation emerges both within and between major lineages [34]. Table 1 displays simulation features. The selection column designates the reaction(s) on which selective pressure operates. The following two columns specify the number of generations spent equilibrating and branching. The phylogenies, all downloaded as Newick files from TimeTree with branch lengths measures in millions of years [35], were selected based on potential relevance to the metabolic model (e.g., a yeast phylogeny for yeast metabolism).

**Table 1.**
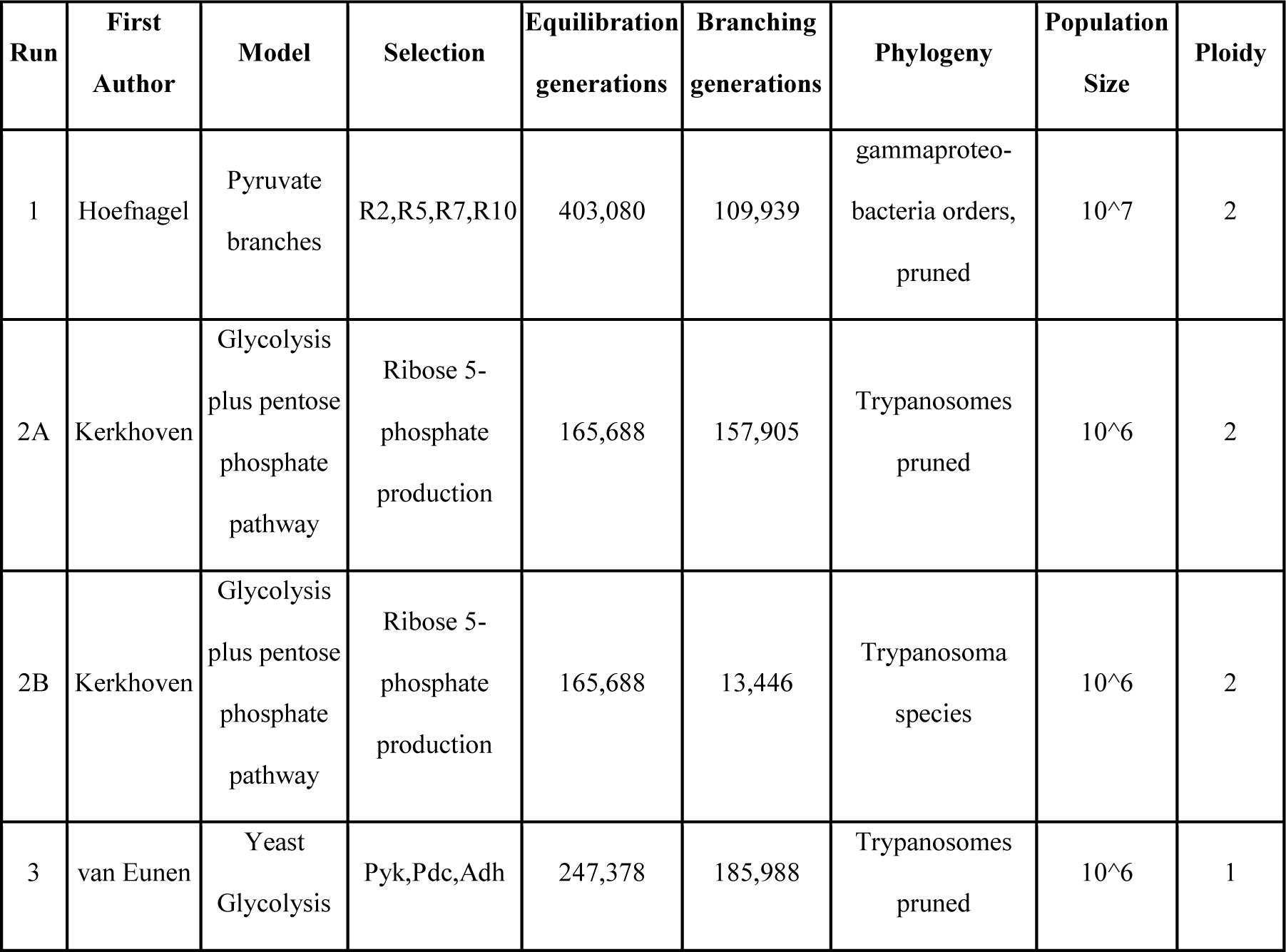
Information about each run, which includes equilibration and branching simulations. Runs 2A and 2B share the same equilibration.

| Run | First Author | Model | Selection | Equilibration generations | Branching generations | Phylogeny | Population Size | Ploidy |
| --- | --- | --- | --- | --- | --- | --- | --- | --- |
| 1 | Hoefnagel | Pyruvate branches | R2,R5,R7,R10 | 403,080 | 109,939 | gammaproteo-bacteria orders, pruned | 10<sup>7</sup> | 2 |
| 2A | Kerkhoven | Glycolysis plus pentose phosphate pathway | Ribose 5-phosphate production | 165,688 | 157,905 | Trypanosomes pruned | 10<sup>6</sup> | 2 |
| 2B | Kerkhoven | Glycolysis plus pentose phosphate pathway | Ribose 5-phosphate production | 165,688 | 13,446 | Trypanosoma species | 10<sup>6</sup> | 2 |
| 3 | van Eunen | Yeast Glycolysis | Pyk,Pdc,Adh | 247,378 | 185,988 | Trypanosomes pruned | 10<sup>6</sup> | 1 |

Run 1 had equally weighted selection on fluxes of the reactions synthesizing outputs of the model network: R2: L-lactate dehydrogenase; R5: acetate kinase; R7: alcohol dehydrogenase; R10: acetoin efflux; R11: acetoin dehydrogenase. Figure 2 shows the population fitnesses during the equilibration process. When equilibrium autodetection does not meet its criteria by 1 million generations, the program begins branching simulation. If desired, equilibration can be extended by the user, but this practical threshold has during development been sufficient for eliminating intolerable levels of systematic directional movement of model parameters in the subsequent phylogenetic simulation. Figure 3 shows the probabilities of fixation for all mutations proposed during equilibration. As most mutations should result in either slightly or substantially deleterious change, modes toward 0 and under 0.5 are expected in a frequency distribution. The mode at 0.5 represents mutations to model parameters that have no effect on fluxes under selection.

**Figure 2.**
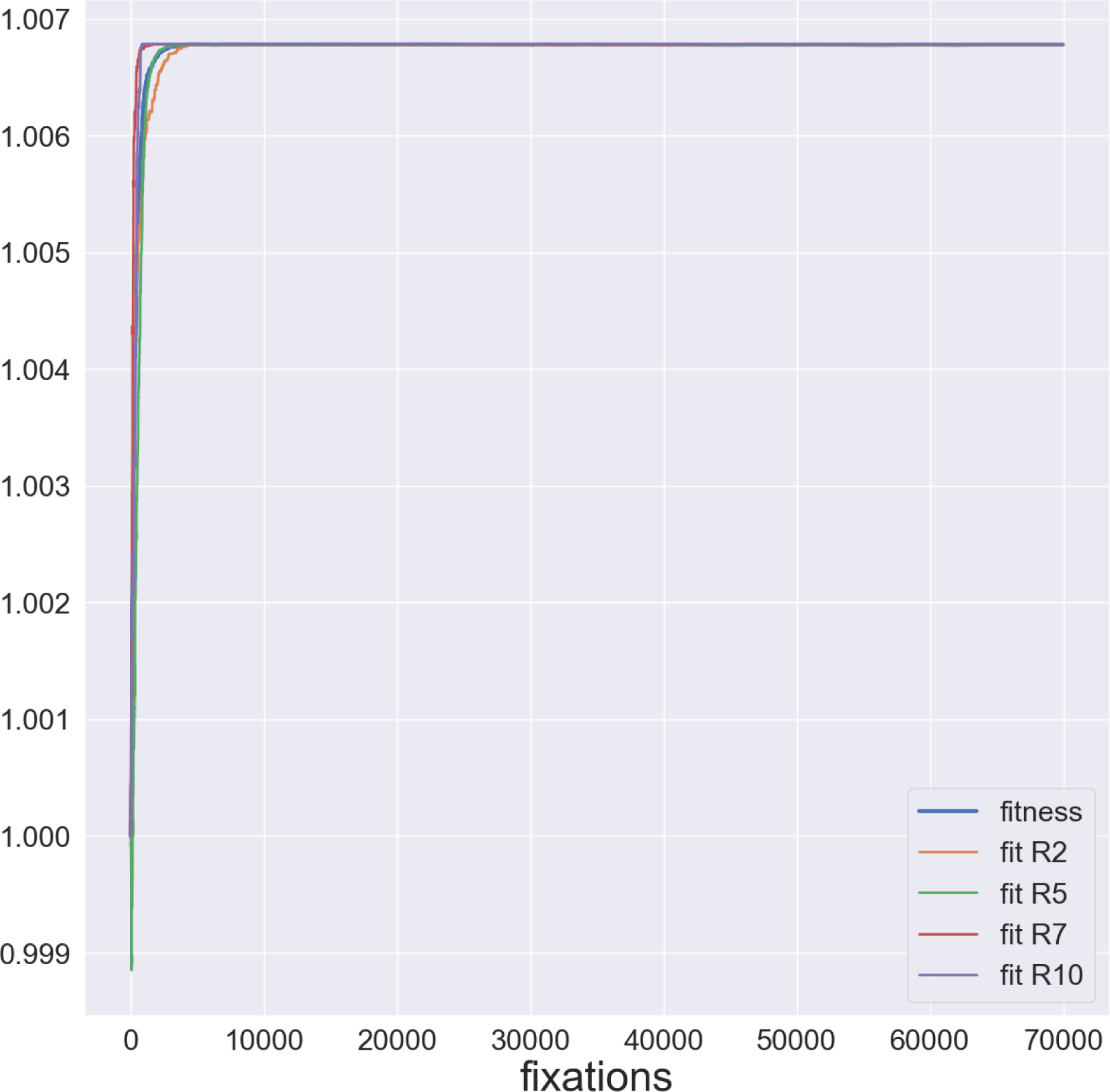
Fitnesses during equilibration in Run 1 (Hoefnagel). Weighted (population) fitness is the product of the fitnesses, equally weighted, of each flux under selection. (R2: L-lactate dehydrogenase; R5: acetate kinase; R7: alcohol dehydrogenase; R10: acetoin efflux)

**Figure 3.**
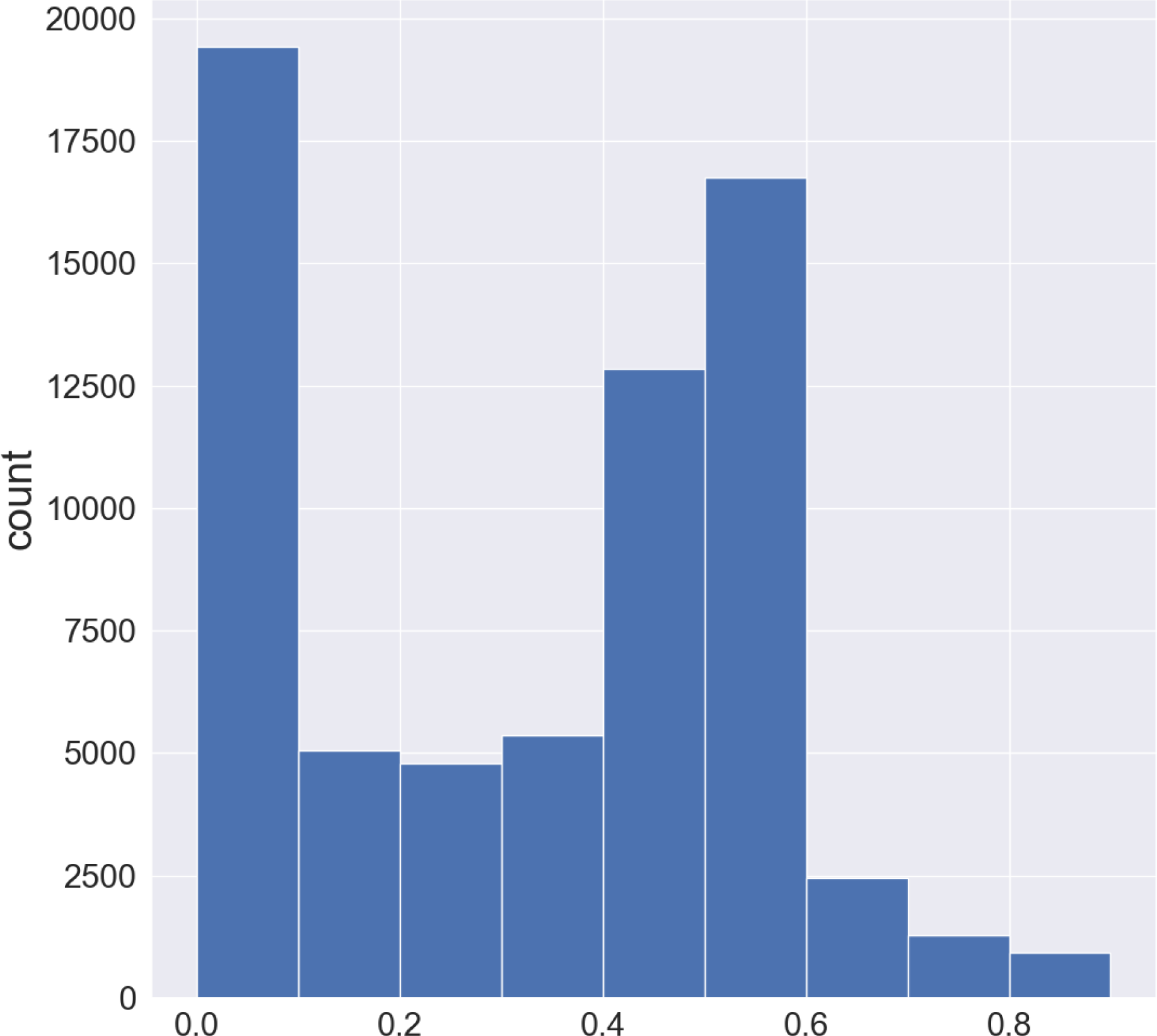
Frequency distribution of probabilities of fixation for all mutations (fixed and unfixed) during equilibration in Run 1.

The input to the following two networks is extracellular glucose, held at a constant value derived from the initial condition described in the model. Simulation 2 ran with selection on ribose-5 phosphate production. It is expected that models can reach steady state, but not that they are imported at evolutionary equilibrium. Under the mutational and selective scheme, most parameter values will show systematic directional movement until a stable balance is achieved, at which point it is most sensible to begin branching the population through the phylogeny. For Simulations 2A&B, the same equilibrated root population was used as the starting point for both branching simulations. Simulation 2A ran on a pruned trypanosome species tree to demonstrate the program’s automatic branch length determination, while 2B ran on the unpruned tree, but for only 1% of the automatically determined generations. It was not practical to fully simulate a large tree with available computational resources. Figure 4 displays lineage-specific fitness for these simulations. Most fluxes, initial concentrations, and non-binding constant parameters reached steady values like the population fitness.

**Figure 4.**
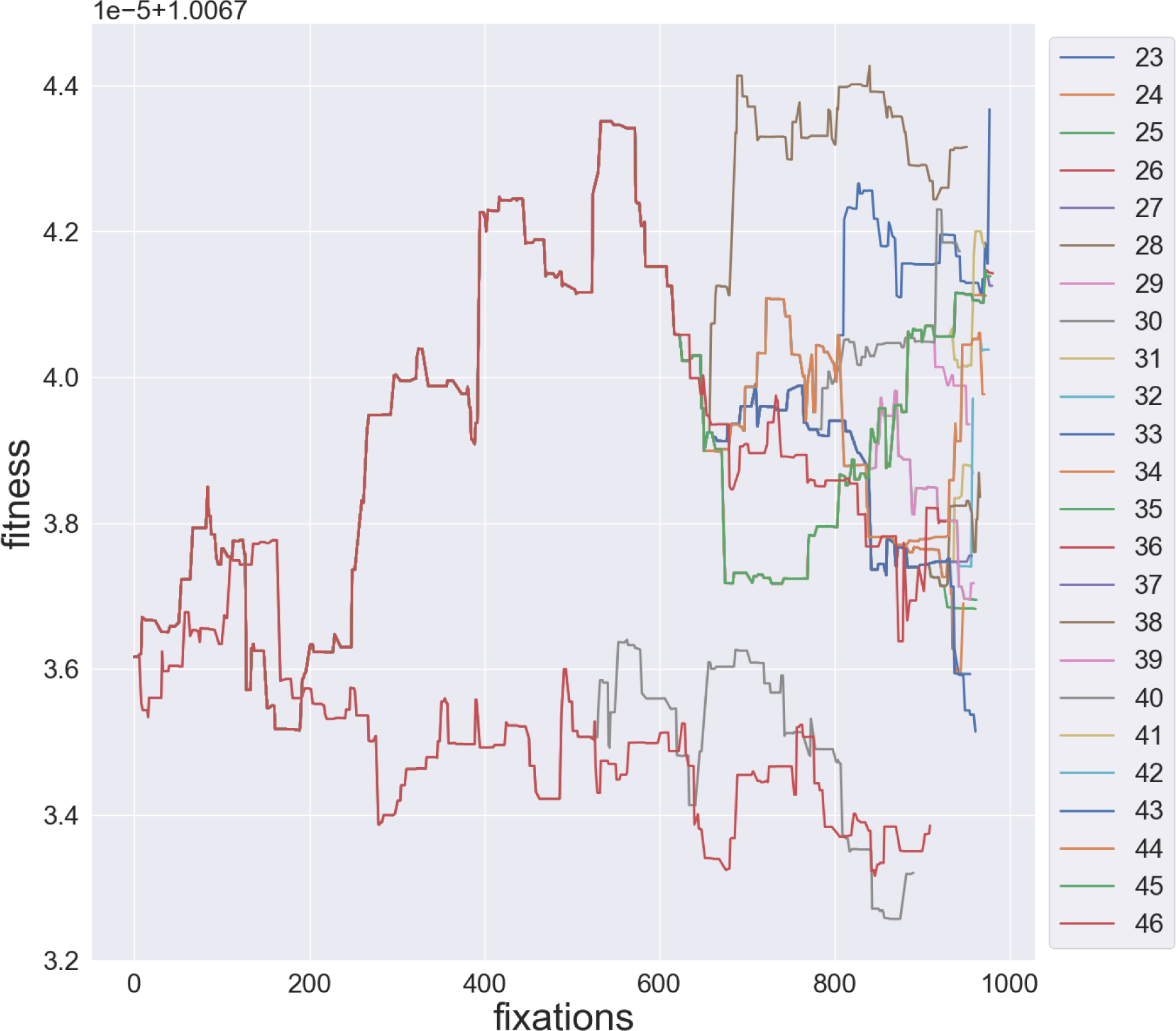
Population fitness along each lineage in Run 2B.

Figure 5 presents example model parameters from Simulation 2A. The upper panels display an initial concentration (panel A: dihydroxyacetone phosphate initial concentration) and reaction flux (panel B: ribose-5 phosphate), whose values depend on steady state calculation and so fluctuate with almost every mutation. The lower panels display parameters which themselves mutate, one (panel C: inhibitory binding constant for ATP in aldolase reaction) with systematic upward movement expected of binding constants [2], and another with nonsystematic change (multiple lineages end up above and below the equilibrated value, panel D: binding constant for ATP in hexokinase reaction).

**Figure 5.**
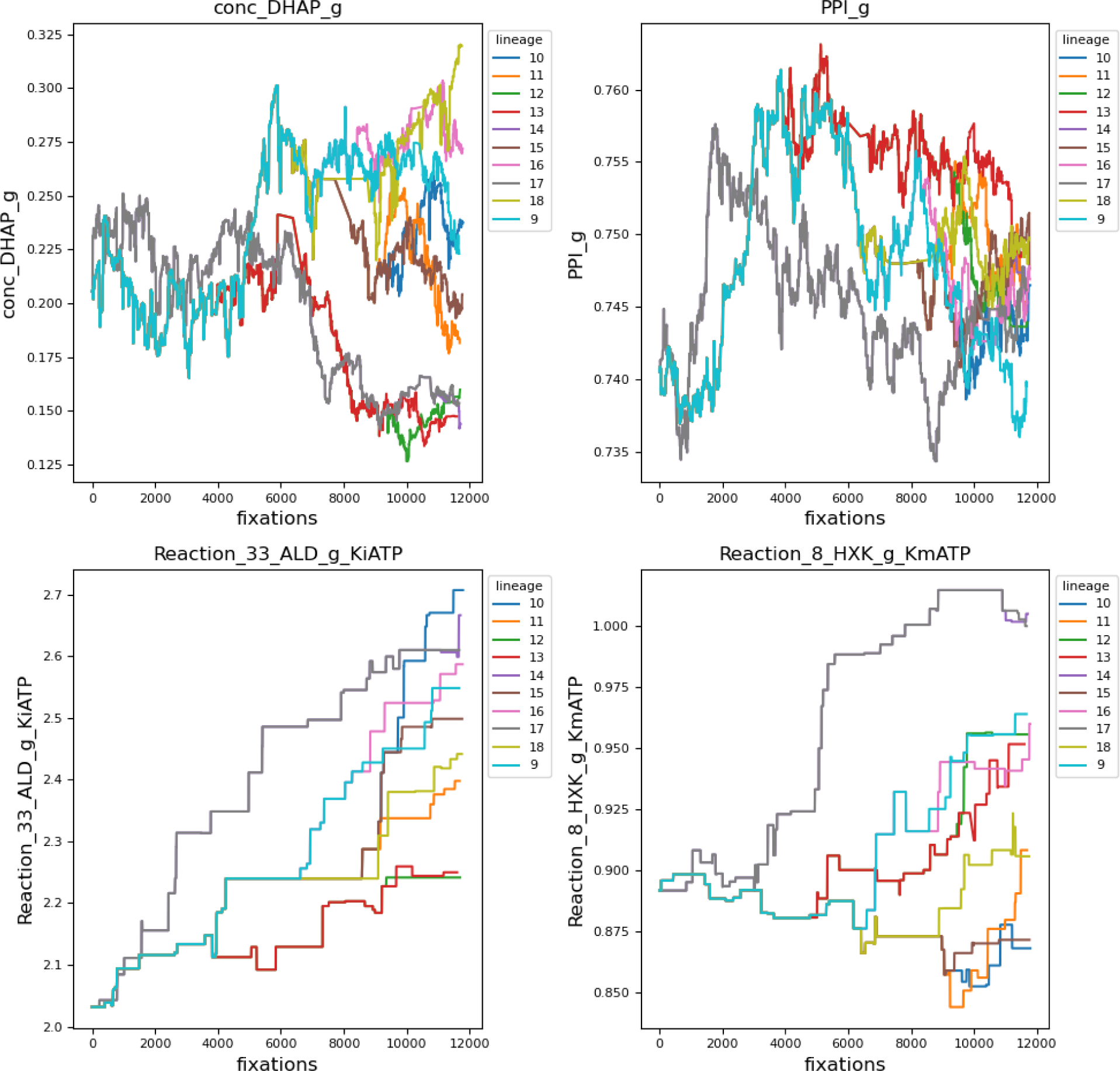
Example parameter evolution along lineages from Simulation 2A. A: dihydroxyacetone phosphate initial concentration. B: ribose-5 phosphate flux. C: inhibitory binding constant for ATP in aldolase reaction. D: binding constant for ATP in hexokinase reaction.

For each parameter of each lineage, the percent difference is calculated between the value at the end of equilibration and at the end of branching. These percentages, for all heatmaps, are normalized (linearly, with respect to maximum column magnitude) such that positive values fall within the range [0,1] and negative values within [-1,0]. Figures 6 and 7 display the normalized percent changes across lineages for Simulations 2B and 3 respectively. When multiple branching simulations are run from the same root over the same phylogeny, the program will average percent changes across them. Figure 8 shows the average percent change across 3 branching simulations for Simulation 1, running at 1% of the automatically determined number of generations.

**Figure 6.**
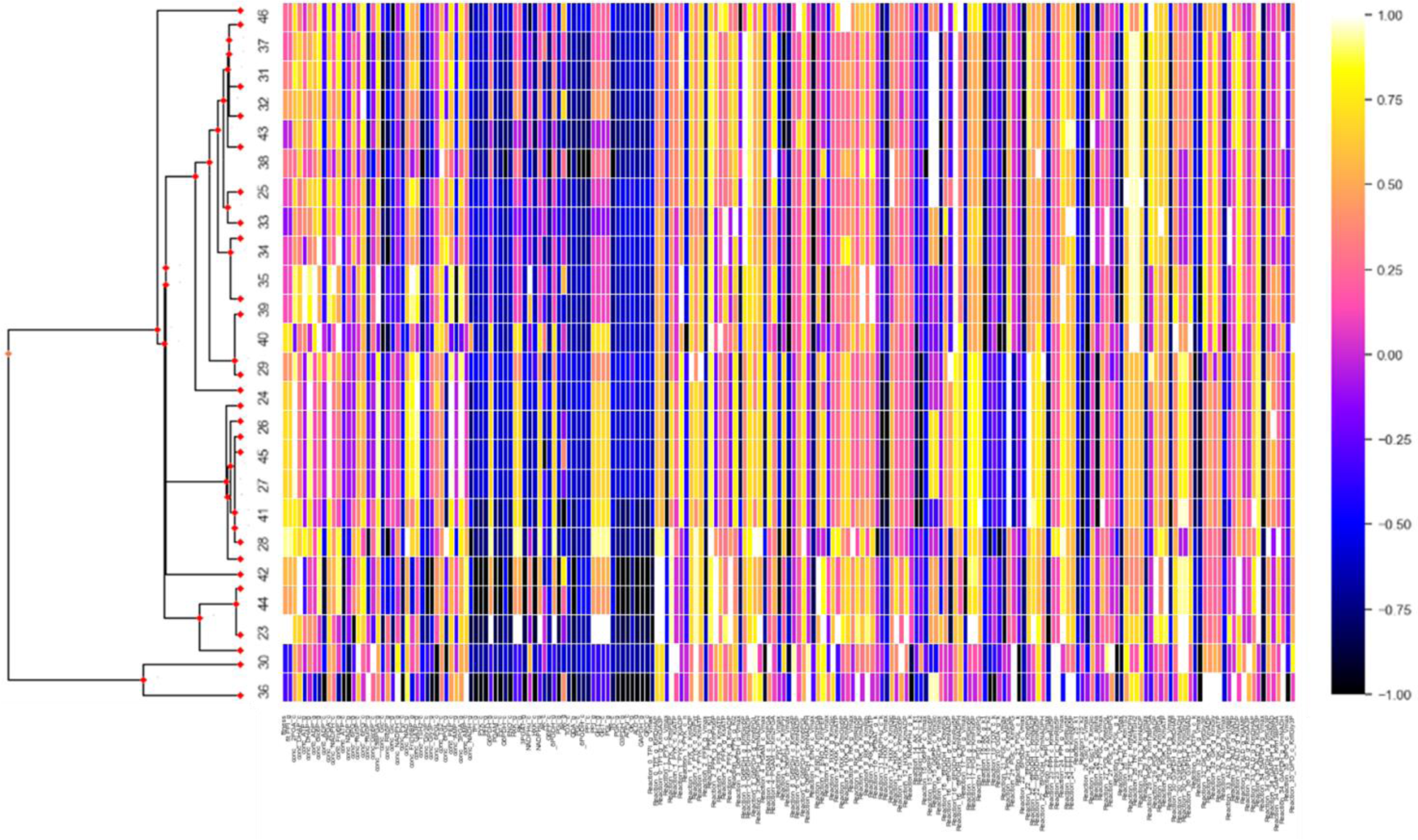
Normalized percent change from initial to final values for lineages in Simulation 2B. Columns represent all model values (fitnesses, fluxes, initial concentrations, parameters) while rows represent external nodes in the Trypanosoma species phylogeny (left).

**Figure 7.**
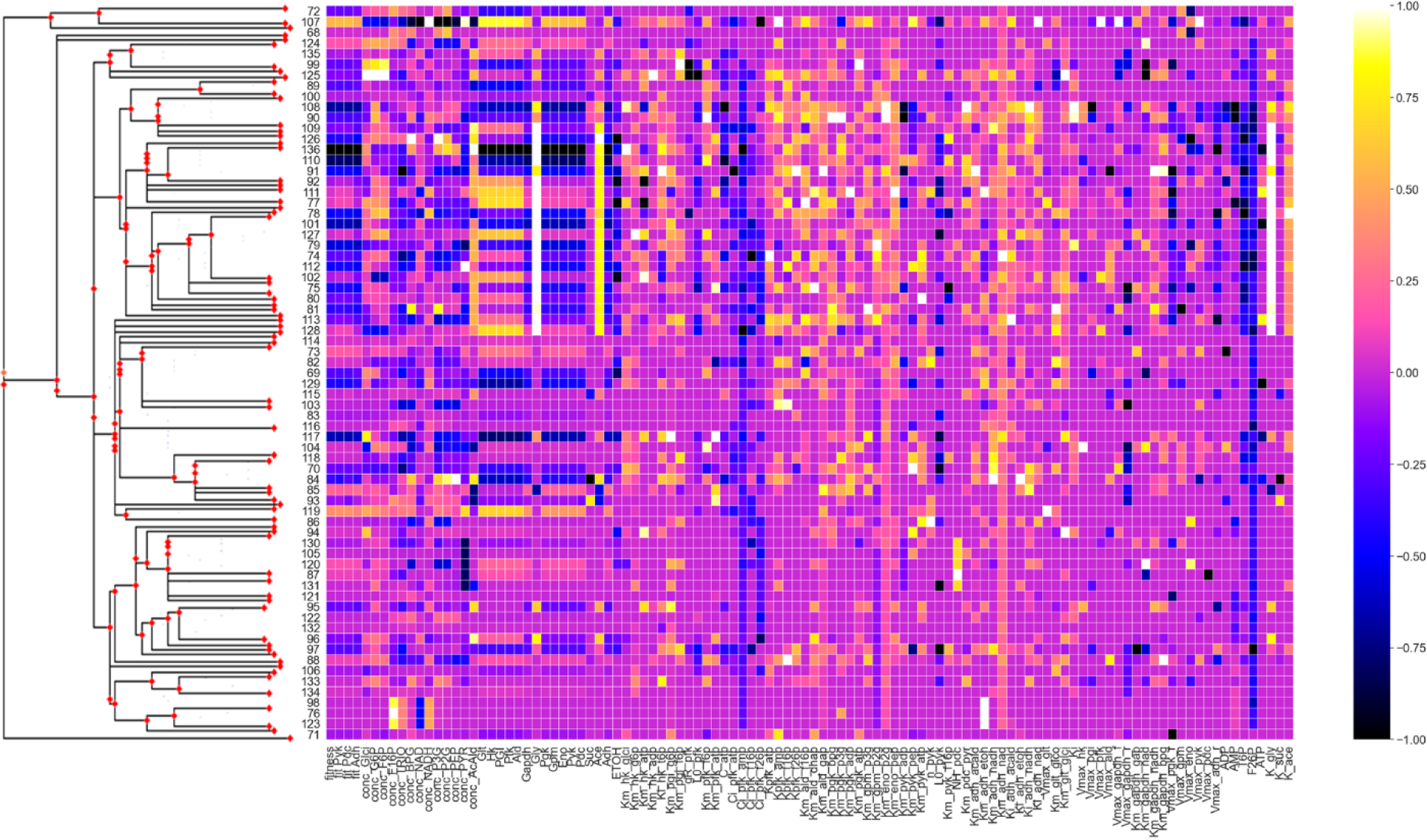
Normalized percent change from initial to final values for lineages in Run 3. Pruned Trypanosoma species phylogeny on the left.

**Figure 8.**
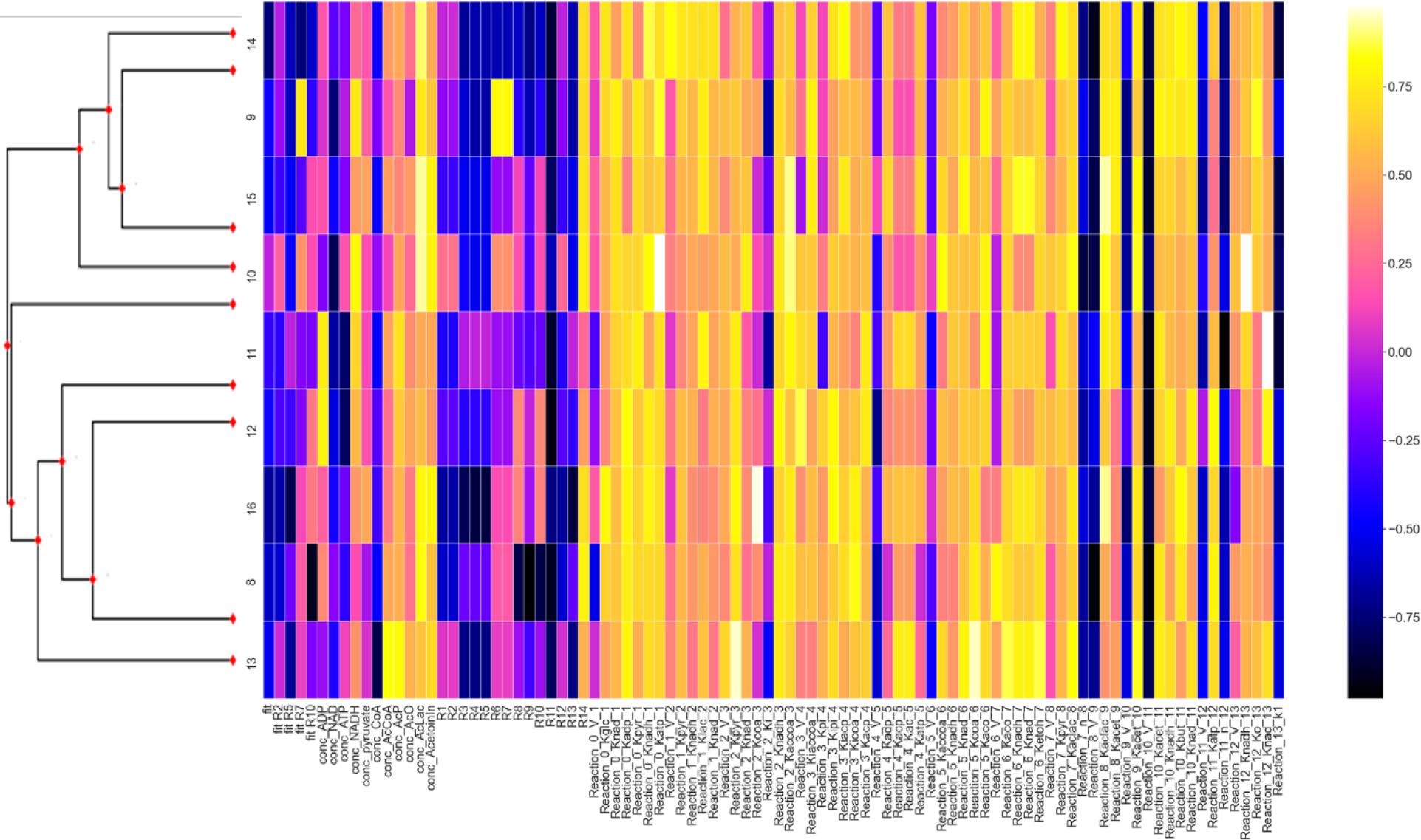
Average normalized percent change for each lineage across 3 branching simulations for Run 1 with pruned gammaproteobacterial order phylogeny on the left.

It is clear in the heatmap that some parameters are moving in unison in parallel across lineages while other parameters move in opposite directions. Those that move in unison are presumably still out of equilibrium from the starting conditions, reflecting rare values among all of the potential values underlying high fitness space. Those that move in opposite directions are a more biologically interesting phenomenon and reflect the alternatives in genotype space corresponding with phenotype and fitness spaces that might be differentially observed as solutions in different genomes from extant species.

## Discussion

Studying the evolution of one gene in isolation can elide the interplay of mutation, selection, drift, and compensatory changes, as well as the resulting equilibrium. While it is commonly assumed that a mutational effect acts independently of genetic background, these simulations contextualize all genetic variants in a network and investigate their effect on pathway flux and population fitness [2]. The PEMPS software simulates enzyme co-evolutionary dynamics under selection on an input (branching) metabolic pathway flux over an input phylogeny. This model could help identify selection on metabolic pathways as well as infer ancestral characteristics.

The relative importance of coding sequence vs. gene expression alterations has been a point of debate in molecular evolution. A Michaelis-Menten kinetics-based mechanistic model, such as those simulated here, if integrated with a protein-level mutational model, could potentially help investigate the phenotypic effects of changes in the level of gene expression (i.e., protein concentration) and functional changes to protein sequence that affect reaction parameters like the binding and catalytic constants. They are naturally intertwined in this framework. Both adaptive and compensatory changes in molecular evolution could be elucidated in studies on mutational opportunity [2].

The limitations to this project include the fact that the metabolic network is fixed, and not temporally regulated, which fails to map onto the dynamic reality that in many cases may much better represent metabolic pathway evolution. Population sizes are assumed to be constant along a phylogenetic tree branch. There is no treatment of heterozygosity in diploid genomes, with the absence of a population model. Furthermore, runtime is cumbersome: branching for even small trees can take multiple days on a typical PC. Because it is optimal to run multiple simulations, copious time must be allocated to generate the data. Finally, while PEMPS can incorporate enzyme kinetic parameters and rate laws from empirical models of steady state flux, these models require extensive experimentation to generate, so data availability remains far from ideal.

It should be noted that network’s architecture is not naturally permanently fixed, which is an implicit assumption of this model. Gene duplication, along with reactions involved in other networks, may alter the dynamics in ways not elucidated by modeling a static pathway structure [36, 37]. Furthermore, many pathways are temporally regulated, so selection does not act on constant flux [2]. The dynamics resulting from modeling such a system could be different than those observed here, which implement no temporal regulation [2]. Pathway parameters involving concentrations may be different in different cell types [38].

While kinetic models achieve the highest level of detail among descriptions of metabolic networks [3], they require that a multitude of parameters and values be empirically determined. Because this is often very difficult, kinetic modeling generally suffers from a shortage of information. Simplifying the kinetic equations without sacrificing similarity to in vivo calculations can address this issue. One method for a simplified kinetic description of metabolic pathways is biochemical systems theory, which focuses on a unified power-law-based treatment of nonlinear rate formulas [39]. A synergistic system of differential equations can be determined from rate constants, metabolite concentrations, and kinetic orders. Biochemical systems theory, however, depends on an accurately defined reference state [3], which may not always be possible to ascertain. Another approach to simplifying the mathematical description of metabolic networks is the linear-logarithmic kinetics (lin-log) framework [40], in which fluxes are formulated based on their deviation from a defined reference state containing the reaction rate, metabolite concentration, and total enzyme activity. This approach can result in better performance than power-law approximations because the kinetic orders are not held constant but vary with metabolite concentrations [3].

Sets of first-order, coupled ODEs have traditionally been used to calculate the time evolution of chemical reactions with the assumptions that the system is homogeneous and that it is suitable to model change in molecular populations as continuous and deterministic processes. However, this latter assumption is usually inappropriate for chemical systems smaller than a test tube, because molecules change in integer quantities and are subject to stochastic effects (knowledge of the current molecular profile in a system is insufficient to predict certainly the future composition) [41]. While arguments from kinetic theory favor the stochastic formulation as more realistic, its reliance on a single differential-difference (“master”) equation can result in mathematical intractability [42], perhaps explaining why all systems encountered in BioModels during development use sets of ODEs.

Improvements to kinetic modeling could result from increasing resolution of metabolic data, standardizing data and modeling steps, and connecting models with other regulatory systems. Despite the capacity to generate metabolic pathways from genomic data, construction of a kinetic model from such a pathway remains a manual process complexified by the inconsistent availability of necessary experimental data on reaction values and parameters [3]. The above approaches to compensating for patchy experimental data, as well as the general improvements, could lead to interesting future studies, including expansions to the current project. The future directions for this study include incorporating time-dependent dynamics in enzyme flux, gene duplication events along the phylogeny, and a feature to allow the user to enter arbitrary pathways.

## Conclusion

Using a mutation-selection modeling framework with fitness derived from ODE-based steady-state flux calculations, the PEMPS program runs on Linux to simulate evolution of metabolic pathways over a phylogeny. The user provides an SBML-formatted kinetic model, a Newick-formatted phylogenetic tree, and a variety of specifications including population size(s), ploidy, and the set of reaction fluxes to subject to selective pressure. Both compensatory and adaptive fixations can emerge in such models and indicate single nucleotide polymorphisms and fixed differences with potential effects on phenotype. If used for inference, this model could ultimately enable detection of selection on metabolic pathways as well as inference of ancestral states for metabolic pathway function. Raw data along with graphical depictions are generated in the output file, and have been consistent with *a priori* expectations of fixation probabilities and systematic change in model parameters.

## Availability and requirements

Project name: PEMPS

Project home page: https://github.com/nmccloskey/PEMPS

Operating system(s): Linux Programming language: Python

Other requirements: Python version 3.8.10 License: Mozilla Public License 2.0

Any restrictions to use by non-academics: None

## List of Abbreviations

PEMPS: Phylogenetic Evolution of Metabolic Pathway Simulator

COPASI: Complex Pathway Simulator

Km: Michaelis binding constant

Keq: Equilibrium constant in Haldane’s relationship

Vmax: Enzyme maximum velocity

## Declarations

### Ethics approval and consent to participate

Not applicable

### Consent for publication

Not applicable

## Availability of data and materials

Software available at https://github.com/nmccloskey/PEMPS.

## Authors’ contributions

DAL conceived the idea for the project. NSM wrote the code. AM helped test the program. NSM wrote the first version of the manuscript, with substantial writing contributions from DAL. All authors approved the final version of the manuscript.

## Acknowledgements

We thank other members of the Liberles Group for helpful comments.

## Additional Files

None

